# Oocyte Aging is Controlled by Mitogen Activated Protein Kinase Signaling

**DOI:** 10.1101/2020.10.29.360693

**Authors:** Hanna Achache, Roni Falk, Noam Lerner, Tsevi Beatus, Yonatan B. Tzur

**Affiliations:** Department of Genetics, Institute of Life Sciences, The Hebrew University of Jerusalem, Jerusalem 91904, Israel; Department of Neurobiology, The Institute of Life Science, The Hebrew University of Jerusalem, Jerusalem 91904, Israel; The Alexander Grass Center for Bioengineering, The Rachel and Selim Benin School of Computer Science and Engineering, The Hebrew University of Jerusalem, Jerusalem 91904, Israel

## Abstract

Oogenesis is one of the first processes to fail during aging. In women, most oocytes cannot successfully complete meiotic divisions during the fourth decade of life. Studies of the nematode *Caenorhabditis elegans* have uncovered conserved genetic pathways that control lifespan, but our knowledge regarding reproductive aging in worms and humans is limited. Specifically, little is known about germline internal signals that dictate the oogonial biological clock. Here, we report a thorough characterization of the changes in the worm germline during aging. We found that shortly after ovulation halts, germline proliferation declines, while apoptosis continues, leading to a gradual reduction in germ-cell numbers. In late aging stages, we observed that meiotic progression is disturbed and crossover designation and DNA double-strand break repair decrease. In addition, we detected a decline in the quality of mature oocytes during aging, as reflected by decreasing size and elongation of interhomolog distance, a phenotype also observed in human oocytes. Many of these altered processes were previously attributed to MAPK signaling variations in young worms. In support of this, we observed changes in activation dynamics of MPK-1 during aging. We therefore tested the hypothesis that MAPK controls oocyte quality in aged worms using both genetic and pharmacological tools. We found that in mutants with high levels of activated MPK-1, oocyte quality deteriorates more rapidly than in wild-type worms, whereas reduction of MPK-1 levels enhances quality. Thus, our data indicate that MAPK signaling controls germline aging and could be used to attenuate the rate of oogenesis quality decline.

## Introduction

Aging leads to a gradual decline and failure of physiological processes. One of the first processes to fail during metazoan aging is oogenesis (Andux & Ellis 2008; Luo *et al.* 2010; Nagaoka *et al.* 2012; Webster & Schuh 2017; Greenblatt *et al.* 2019; Gruhn *et al.* 2019). In women, oocytes enter meiosis during maternal embryogenesis, arrest at the end of meiotic prophase I, and remain quiescent for decades. Over time, the quality and quantity of these oocytes decreases, concurrently with reduced fertility and increased occurrence of aneuploidy (te Velde & Pearson 2002; Bentov *et al.* 2011; Eichenlaub-Ritter *et al.* 2011; Duncan *et al.* 2012; Lord & Aitken 2013). Knowledge of the genetic and molecular mechanisms that govern this aging process is currently limited.

Studies in the nematode *Caenorhabditis elegans* have played pivotal roles in our understanding of the genetic contribution to longevity and aging. The *C. elegans* system has the advantages of short lifespan (2–3 weeks), simple genetic setup, and the evolutionary conservation of the longevity pathways (Kenyon 2010; Luo *et al.* 2010; Wilkinson *et al.* 2012). Several inherent properties make *C. elegans* highly suitable for the study of germline aging (Hughes *et al.* 2007; Andux & Ellis 2008; Luo *et al.* 2009). First, oogenesis is continuous and the nuclei in the adult gonad are ordered in a spatio-temporal manner from the germ line stem cells to the mature oocyte (Crittenden *et al.* 1994; Lui & Colaiacovo 2013; Pazdernik & Schedl 2013; Hillers *et al.* 2017). Second, in hermaphrodites, ovulation is continuous as long as self-sperm is available. Once sperm is depleted, oocytes arrest at the end of meiotic prophase I. The hermaphrodite worm remains fertile for several more days and can resume ovulation and fertilization upon mating with males as a response to the introduction of allosperm into the uterus (Hodgkin 1983; Hughes *et al.* 2007; Andux & Ellis 2008; Mendenhall *et al.* 2011; Chasnov 2013; Pickett *et al.* 2013; Kocsisova *et al.* 2019). Thus, unlike human oocytes, worm oocytes are continuously produced because the germ-cell population is proliferative (Crittenden *et al.* 2006; Crittenden & Kimble 2008). Finally, both human and *C. elegans* females reproduce for about one-third of their lifespan (Hughes *et al.* 2007) and thus undergo reproductive aging on proportional time scales.

Building upon these properties, several previous works described different aspects of oogenesis at some phases of germline aging in *C. elegans* (Hughes *et al.* 2007; Andux & Ellis 2008; Luo *et al.* 2009; Luo *et al.* 2010; Ye & Bhalla 2011; Wang *et al.* 2014; Bohnert & Kenyon 2017; Templeman & Murphy 2018). These works showed that aging oocytes undergo gross morphological and functional changes. Morphological changes include the presence of small stacked oocytes and endomitotic nuclei in the proximal gonad (de la Guardia *et al.* 2016; Kocsisova *et al.* 2019). Functional defects that occur during aging involve reduced embryo hatching and stress resistance, decreased oocyte fertilizability, altered crossover distribution, and high incidence of males (Lim *et al.* 2008; Luo *et al.* 2010; Perez *et al.* 2017). In addition, mutations in several genetic pathways extend the fertility term (Luo *et al.* 2010; Hughes *et al.* 2011). Among these, mutations in genes encoding factors involved in the insulin/IGF-1 signaling (IIS), the TGF-ß-Sma/Mab, and the dietary restriction pathways extend the fertility period in worms from both self and allosperm (Reviewed in (Lopez-Otin *et al.* 2013)). Nevertheless, to date no systematic work has analyzed the dynamics of major meiotic processes along all steps of germline aging. Moreover, the germline signals that lead to the specific oogonial changes during normal aging are still largely unknown.

Signals that control developmental processes are often also involved in aging (Blagosklonny & Hall 2009; Gruber *et al.* 2016; Slack 2017). We therefore hypothesized that some signaling pathways that are activated during oogenesis also influence germline and oocyte aging. The MAPK pathway controls oogenesis progression in *C. elegans* (Lee *et al.* 2007; Kim *et al.* 2013; Nadarajan *et al.* 2016). Several proteins that promote or restrict the activation of its terminal kinase, MPK-1, the worm homolog of ERK, and this, in turn, leads to multiple transcriptional and post-transcriptional cellular changes that drive oogonial processes (Church *et al.* 1995; Lackner & Kim 1998; Kritikou *et al.* 2006; Leacock & Reinke 2006; Lee *et al.* 2007; Arur *et al.* 2011; Yin *et al.* 2016; Achache *et al.* 2019).

We found that meiotic progression is altered and processes such as double-strand break repair and crossover designation are reduced in the *C. elegans* germline during aging. These alterations occur concomitantly with a change in spatial activation of MPK-1. During aging, oocyte quality was inversely correlated with and dependent on the level of MAPK activation. Furthermore, in mutants with high levels of activated MPK-1, oocyte quality deteriorated more rapidly than in wild-type worms, whereas reduction of MPK-1 levels enhanced quality. We conclude that MAPK signaling in mature oocytes controls reproductive aging by influencing oocyte and germline quality.

## Results

### Germline aging leads to a reduction in germ cell numbers and altered meiotic staging

*C. elegans* hermaphrodite worms transiently produce sperm during the L3 larval stage and switch to oogenesis in the fourth larval stage (L4) (reviewed in (Schedl 1997)). Oocytes start to be fertilized at the adult stage, and hermaphrodites continue to ovulate until most of the self-sperm is depleted, at which point the oocytes arrest and age (Kim *et al.* 2013; Templeman & Murphy 2018). To study germline aging, we chose to use the N2 wild-type strain instead of feminized mutants as was done previously (e.g., (Hughes *et al.* 2007; Andux & Ellis 2008; Lim *et al.* 2008; Luo *et al.* 2010; de la Guardia *et al.* 2016; Bohnert & Kenyon 2017; Templeman & Murphy 2018)). Our strategy ensured that the aging effects we detected were unrelated to any mutation. We defined four points during the aging process: the onset of reproduction, the beginning of the arrest, the end of the reproductive term by male cross-fertilization, and after the reproductive term (Hughes *et al.* 2007), which can also be described as young, mature, old, and menopausal, respectively. Previous work has shown that most self-progeny of wild-type worms are laid at the second day post L4 and that almost all the embryos are laid within three days (Hughes *et al.* 2007; Pickett *et al.* 2013; Wang *et al.* 2014). We verified that this occurred under our experimental conditions (Fig. S1). A negligible number of oocytes were laid after three days post L4 (under 0.8 on average per worm), and no viable embryo was laid after the fifth day (Fig. S1). Thus, we chose to compare worms on the first (young), fourth (mature), eighth (old), and tenth (menopausal) days after L4 stage.

To find how aging affects germ cell number and developmental stages, we examined dissected gonads stained with DAPI. The *C. elegans* gonad is comprised of two U-shaped arms with nuclei arranged in spatial-developmental order. The proliferative zone is located at the distal end of each arm, and mature oocytes and the spermatheca are found at the proximal end. Nuclei in the proliferative zone undergo mitotic cell cycles to maintain a population of progenitors that enter meiosis in the leptotene/zygotene (LZ, transition) zone. From there, nuclei progress through pachytene, where recombination intermediates mature into crossovers within paired and synapsed homologs. Pachytene nuclei move into diplotene, and finally oocytes mature and cellularize in diakinesis, where six discrete bivalents can be visualized (Fig. 1A). We counted the total number of nuclei in DAPI-stained gonads and observed an overall decrease with age (Fig. 1B). The reduction in germ cell numbers could be due to either a relative reduction in numbers at all meiotic stages or to numbers at specific stages. To determine which is the case in the aging gonad, we quantified the numbers of nuclei at different meiotic stages. The number of LZ nuclei quickly dropped and were reduced at the onset of oocytes arrest (day 1: 108±17; day 4: 13±9; Fig. 1B). We also detected a reduction in the number of pachytene nuclei during aging (Fig. 1B). Thus, germline aging and arrest lead to reductions in germ cell numbers, mostly due to a rapid drop in the number of LZ and pachytene nuclei.

**Figure 1:**
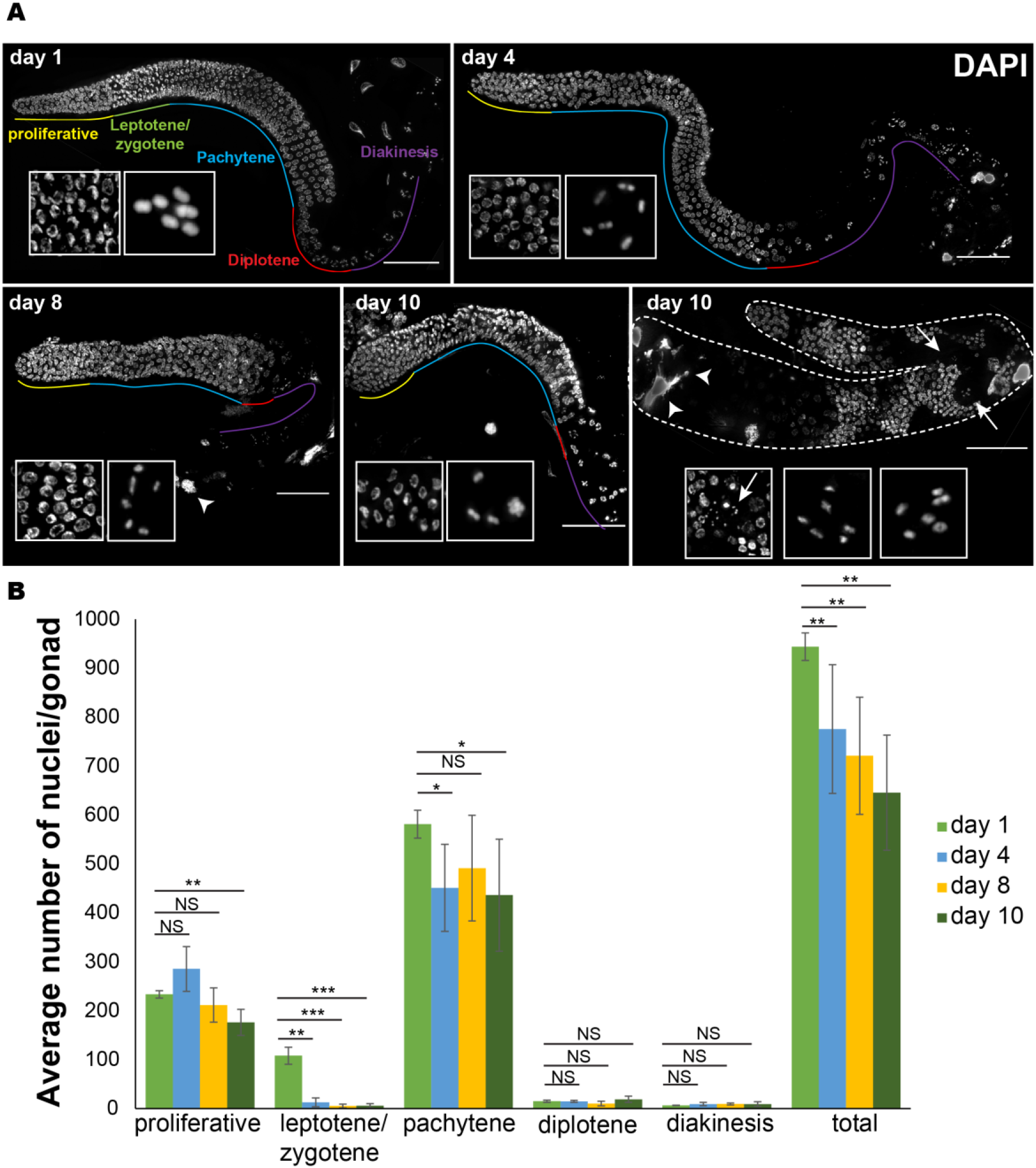
Aging leads to reduced number of germ cells. **A**. Images of DAPI-stained whole mount gonads from worms at the indicated ages. The different oogonial stages are marked. Insets shows early and late stages nuclei. Arrowheads – endomitotic nuclei. Arrows – nuclei with diakinesis morphology located at the distal side. Scale bar = 50 μM. **B**. Average number of total nuclei at different stages per gonad from worms at the indicated ages. Mann-Whitney *p* value: NS – not significant, *<0.05, **<0.01, ***<0.001. N = 6 gonads.

The spatial temporal order in the young adult gonad has been highly advantageous in meiotic studies in this model organism. This order was always present in day 1 and day 4 worms. However, starting at day 8 after the L4 stage, we noticed an increased number of gonads with altered morphology. About 17% of the gonads were very small (with fewer than 500 germ cell nuclei) on day 8, and about 19% were very small on day 10. In addition, there was an increase in gonads with greater than 1000 germ cells (3.5% at day 8 and 18.5% at day 10). In these gonads, the meiotic order was disrupted, and diakinetic-like nuclei with large cytosolic volumes, were observed along the middle of the gonad, and pachytene-like nuclei were observed proximally (Fig. 1A). The mixture of stages indicates that during aging, there is loss of meiotic progression control. A similar phenotype was reported previously in mutants of *kin-18* (Yin *et al.* 2016), an activator of MAPK. Taken together these analyses suggest that germline aging leads first to a reduction in the number of germ cells and then to misregulation of meiotic progression.

### Germ cell proliferation declines with aging

The number of nuclei in the gonad is tightly regulated by a dynamic balance between germ cell proliferation and removal by both oocyte ovulation and apoptotic cell death (Lettre & Hengartner 2006). When sperm are depleted, ovulation ceases almost completely. The reduction in the number of germline nuclei may therefore be due to either an increase in apoptosis or a decrease in mitotically proliferating nuclei. To test the former, we used a strain stably expressing CED-1::GFP, a fusion protein that is expressed in somatic sheath cells, which cluster around each apoptotic corpse during engulfment (Zhou *et al.* 2001; Schumacher *et al.* 2005). This approach is particularly useful for detecting early apoptotic stages. In agreement with a previous publication (de la Guardia *et al.* 2016), in young adult worms we found fewer apoptotic nuclei (4.6±2.1) than in mature worms (8.2±3.6), but the trend was reversed in old worms (4.1±2.4, n=7, Fig. 2A). We were unable to quantify apoptotic levels at day 10, since almost all the transgenic CED-1::GFP worms died before reaching day 10. These results suggest that at the onset of aging, the apoptotic removal of meiocytes increases but then returns to young worm levels.

**Figure 2:**
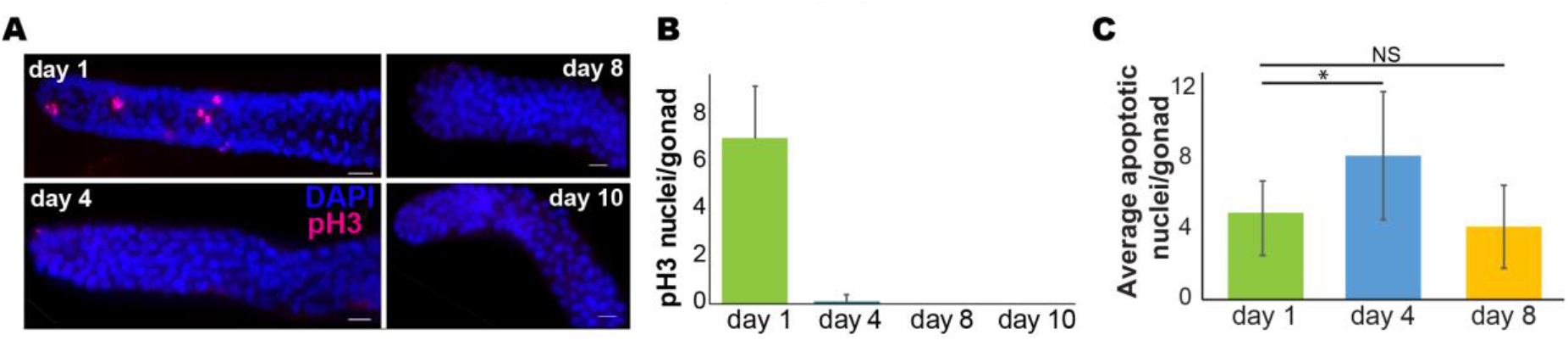
Changes in proliferation and apoptosis. **A**. pH3 (red) and DAPI (blue) staining of proliferative zone from worms at the indicated ages. Scale bar = 10 μM. **B**. Average number of pH3-positive nuclei per gonad. N = 16 gonads for day 1, N = 21 for day 4, N = 10 for day 8, and N = 6 for day 10. **C**. Average number of CED-1::GFP apoptotic nuclei per gonad from worms at the indicated ages. N = 13 gonads. Mann-Whitney *p* value: NS – not significant, *<0.05.

The increase in apoptosis was transient, whereas the reduction in germ cell numbers was continuous. To investigate if proliferation also regulates the overall number of nuclei in the gonad, we monitored the number of germ cells in M phase at days 1, 4, 8 and 10 by staining for a mitosis-specific marker phospho-histone H3 (pH3) (Hans & Dimitrov 2001). We detected a rapid decline in the number of pH3-positive nuclei in the distal region at day 4, and at days 8 and 10 no nuclei were stained with pH3 (Fig. 2B, C). Taken together these results suggest that aging leads to reduction in the number of germ cells due to both reduced proliferation and increased apoptosis.

### Reduced levels of RAD-51 foci in aging gonads

Apoptosis in the worm germline has been correlated with aberrations in DNA double-strand break repair and synapsis (reviewed in (Gartner *et al.* 2008)). To test if the increase in apoptosis is induced by altered dynamics of homologous recombination repair, we quantified the number of repair loci using RAD-51 staining. RAD-51 is a strand exchange protein that has been used extensively to study the DNA double-strand break repair dynamics in the *C. elegans* gonad (Colaiacovo *et al.* 1999; Rinaldo *et al.* 2002; Alpi *et al.* 2003; Bhalla & Dernburg 2005; Hayashi *et al.* 2007; Mets & Meyer 2009; Yu *et al.* 2016). In the young adult worms, the levels of RAD-51 rose following entry into meiotic prophase I and peaked in the early to mid-pachytene stage (Fig. 3A, B), as previously reported (Colaiacovo *et al.* 2003; Achache *et al.* 2019). Numbers of RAD-51 foci were greatly reduced in nuclei at all the stages of meiotic prophase I in aged worms (Fig. 3A, B). Indeed, at day 1, we found an average of 4.7±2.6 foci per nucleus in early pachytene, compared to 1.6±1.3, 1.3±1, and 0.9±0.9 at days 4, 8, and 10, respectively. Thus, after the halt in ovulation, the number of RAD-51 foci drops. This drop in the number of RAD-51 foci might be explained by a progression of the arrested nuclei beyond the removal of RAD-51, together with a reduction in further induction of double-strand breaks. Another option could be a defect ‘ in either the induction or the repair of the DNA double-strand breaks. Interestingly, in gonads of day 8 and 10 nuclei, we detected RAD-51 staining that filled the entire nucleoplasm (Fig. 2A). This type of staining could be the result of fragmented DNA or misregulation of RAD-51 expression. Taken together our results suggest that the increase in apoptosis observed on day 4 is unlikely to be the result of perturbations in the DNA repair mechanism.

**Figure 3:**
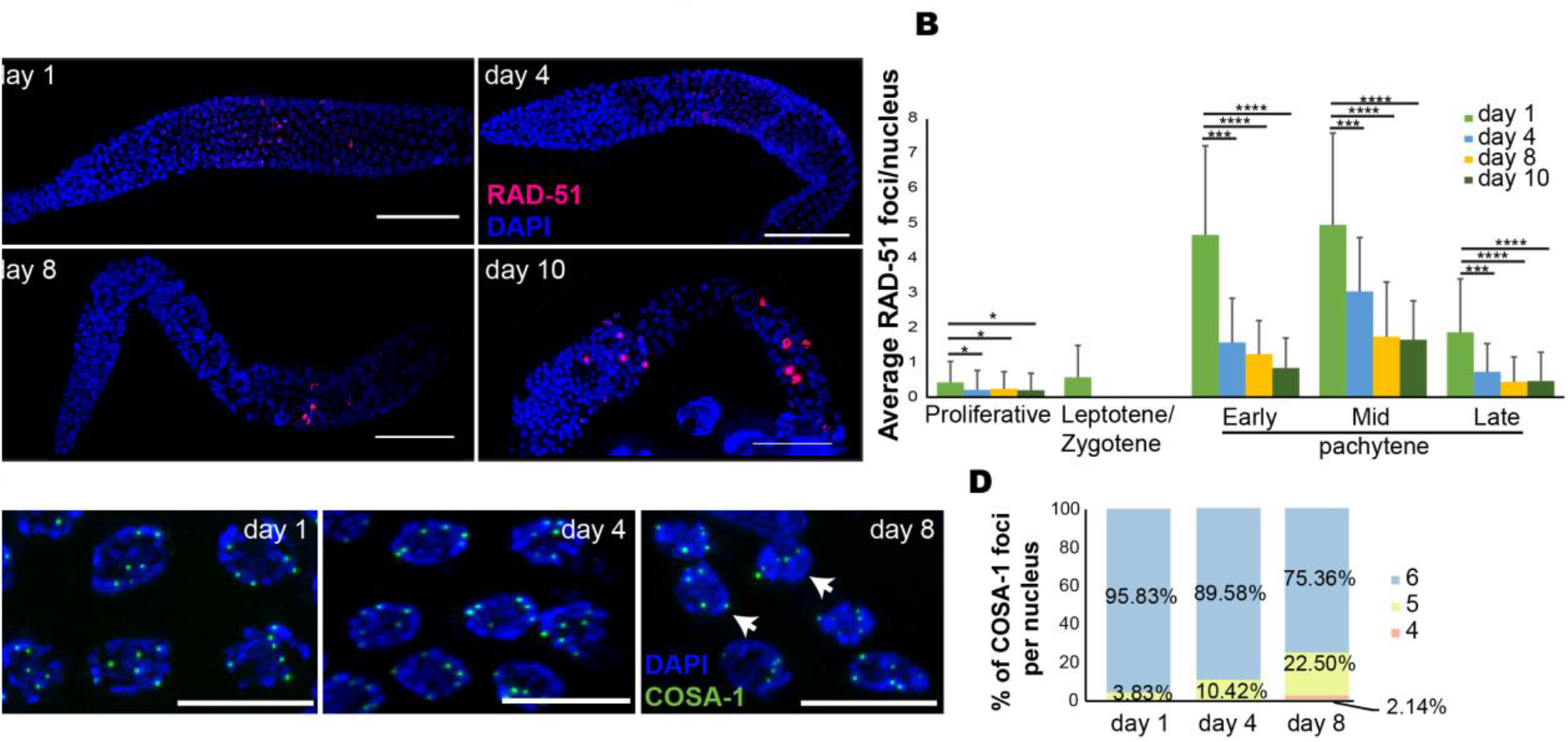
Reduced RAD-51 foci and crossover designation markers in aged germlines. **A**. RAD-51 (red) and DAPI (blue) staining of whole mount gonad arms at the indicated ages. **B**. Average number of RAD-51 foci per gonad nuclei at the different oogonial stages. Scale bar = 50 μM. Mann-Whitney *p* value: NS - not significant, *<0.05, ***<0.001, ****<0.0001. **C.** COSA-1::GFP (green) and DAPI (blue) staining of late pachytene nuclei at the indicated ages. Scale bar = 10 μM. **D**. Histogram of the relative percentages of nuclei with four, five, and six COSA-1 foci per late pachytene nucleus at the different ages. N = 200 nuclei.

### Synaptonemal complex formation is unchanged during germline aging

During meiosis, the formation of a proteinaceous structure known as the synaptonemal complex (SC) stabilizes pairing interactions and promotes the completion of crossover recombination. SCs assemble along the lengths of the paired chromosomes to keep them closely associated and aligned (Colaiacovo *et al.* 2003; Couteau & Zetka 2005; Hayashi *et al.* 2010; Schild-Prufert *et al.* 2011). This zipper-like structure is composed of lateral element proteins that are recruited to the chromosome axes and to central region proteins that localize between them and keep the homologs aligned (reviewed in (Zetka 2009)). Failure to properly form the SC increases apoptosis levels in the gonad (Bohr *et al.* 2016). To test if the decrease in apoptosis levels on day 4 relative to day 1 is correlated with aberrant synapsis, we performed immunostaining using antibodies against SYP-2, a central region protein (Colaiacovo *et al.* 2003), and HTP-3, an axial component of the SC (MacQueen *et al.* 2002; Goodyer *et al.* 2008; Severson *et al.* 2009). In mid-pachytene stages, these proteins co-localize on the DAPI-stained tracks in young adult worms (Fig. S2), indicating proper formation of the SC. We observed similar patterns at all aging timepoints with no indications of loss or partial synapsis (Fig. S2). Thus, at least at the level of this observation, synapsis is not altered with germline aging. Taken together, these results open the possibility that factors other than synapsis and repair of DNA double-strand breaks contribute to the increased apoptosis observed on day 4 after L4 in worm gonads.

### Germline aging leads to small reduction in crossover designation

Interhomolog crossover recombination is dependent on proper repair of DNA double-strand breaks, and reduced crossovers have been suggested to play roles in aneuploidy in advanced aged mothers (reviewed in (Webster & Schuh 2017)). To determine if the reduction in RAD-51 foci numbers with maternal age are accompanied by changes in crossover designation, we examined the loading of GFP-tagged COSA-1 onto meiotic chromosomes. COSA-1 localizes to the single crossover site in each homolog pair in late prophase I (Yokoo *et al.* 2012) and is the earliest known marker for crossovers in *C. elegans.* At day 1, almost all nuclei (96%) in the last five rows of the pachytene stages of the young adult gonad showed six COSA-1 foci corresponding to the six crossovers sites per nucleus present in *C. elegans* (Fig. 3C, D). Aging led to a gradual decrease in crossover designation. On day 4, 10.4% of the nuclei had five foci, and on day 8, 22.5% had only five foci (Fig. 3C, D). Moreover, on day 8 after L4, 2% of the nuclei in late pachytene had only four COSA-1::GFP foci (Fig. 3C, D). Interestingly, a reduction in the number of COSA-1 foci was observed in late prophase I stages in *kin-18* mutants (Yin *et al.* 2016). Similar to the CED-1::GFP worms, almost all the worms of COSA-1::GFP transgenic strain died before day 10, thus we were unable to collect relevant data for that stage. These results suggest that aging leads to reduction in crossover recombination designation.

### The distance between homologous chromosomes increases in old worms

In human, the percentage of meiotic chromosomal mis-segregation exponentially increases with maternal age (Hassold & Hunt 2001; Koehler *et al.* 2006). This has been attributed to the time oocytes are arrested, and, indeed, increased premature dissociation of chromosomes has been observed in oocytes in aging women (Subramanian & Bickel 2008; Lister *et al.* 2010; Tsutsumi *et al.* 2014). Nevertheless, the magnitude of premature dissociations is lower than the aneuploidy rate, suggesting that other factors control the arrested oocyte quality and potential to complete the divisions (Nagaoka *et al.* 2012). The increased levels of oocytes with only five COSA-1::GFP foci in old worms raises the possibility that aging leads to reduced crossovers, which in turn should lead to presence of univalent chromosomes in mature oocytes. In young adult worms, the six bivalents of *C. elegans* are almost always detected as six separate DAPI-stained bodies. When homologous chromosomes either do not undergo crossovers or separate before anaphase I, more than six bodies are expected. We did not observe an increase in extra bodies in aged oocytes (Fig. 1A). In fact, we noticed an increase in the number of oocytes in which the bivalents were in very close proximity, and these chromosomes seemed connected at the resolution level of our microscopy system (12.5% of the mature oocytes contained 5 bivalents at day 10 vs. 3% at day 1, Fig. 1A, N = 32). This suggests that aging does not lead to nuclei with non-crossover chromosomes; however, nuclei with non-crossover chromosomes may be removed by apoptosis.

Although homologs are attached during diakinesis, we hypothesize that this attachment weakens with age. This hypothesis predicts that homologs in aged oocytes are more spatially separated than in young oocytes, as previously observed in mouse and human (Gruhn *et al.* 2019; Zielinska *et al.* 2019). To test this hypothesis, we used a fluorescence microscope to capture the 3D chromatin density of the chromosomes in mature oocytes in both young and menopausal worms (Fig. 4A). To identify individual chromosomes within each pair of homologs, we used an unbiased 3D Gaussian-mixture model with two Gaussians that was fitted to the measured chromatin density. The spatial separation between the homologs was quantified by projecting all images in the z-stack into a single 2D chromatin density map and calculating the 2D distance (*L*) between the projected centers of the two Gaussians (Fig. 4B). This analysis showed that *L* is significantly longer in oocytes of menopausal vs. young worms (Fig. 4C).

**Figure 4:**
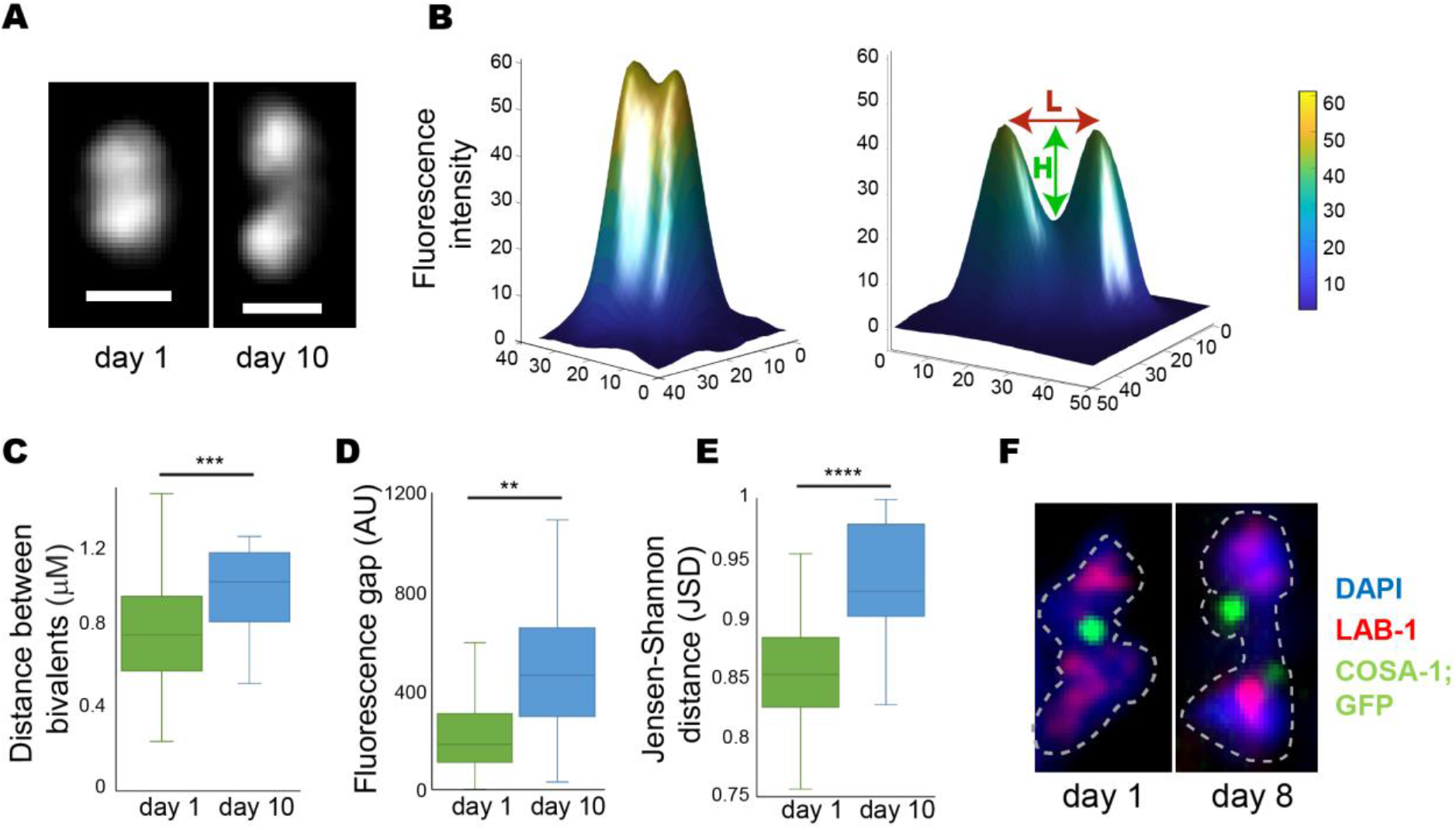
Homologs are located at greater distance from each other in old worms. **A**. The z-projected images of DAPI-stained bivalents in mature oocytes at day 1 and day 10. Scale bar = 1 μM. **B**. 2D chromatin density maps obtained by projecting the z-stacks in panel A for day 1 and day 10 bivalents from mature oocytes. Chromatin density is indicated both by color and surface height. The 2D distance between the projected Gaussian centers is defined as *L.* The difference in fluorescence between the minimum value along *L* and the mean value of the two Gaussians centers is defined as *H.* **C**. Box and whisker plot of the distance *L.* N = 20. **D**. Box and whisker plot of the fluorescence difference *H.* N = 20. **E**. Box and whisker plot of the Jensen-Shannon distance between the 3D Gaussian distributions of the two homologs in each pair. N = 20. **F**. Images of bivalents from day 1 and day 8 mature oocytes stained for LAB-1 (red), COSA-1::GFP (green), and DAPI (blue). N = 32.

If this longer distance is caused by weakening of the sister chromatid cohesion, then less chromatin is expected to be present at the interface between the two homologs (known as the short arms). To test this prediction, we analyzed the fluorescence profile between the two peaks of the projected 2D chromatin map along the line *L.* We define *H* as the fluorescence difference between the minimum value along *L* and the mean height of the two Gaussians centers (Fig. 4D). The value of *H* is significantly greater in menopausal worms than in young worms, suggesting that, indeed, there is less chromatin at the short arms. To further verify these results, we used the Jensen–Shannon divergence to quantify the overlap between the two Gaussian probability distributions in 3D as described (Lin 1991; Endres & Schindelin 2003). We found significantly lower levels of overlap between the homologs in menopausal worms than in young worms (Fig. 4E), supporting the finding that homologs in menopausal worms are more spatially separated than in younger worms.

Interestingly, when we imaged oocytes in aged COSA-1::GFP worms, we noticed that 82.5% of the oocytes contained bivalents with double COSA-1 foci at the chiasmata region (Fig. 4F). Together with the DAPI staining, this observation supports the hypothesis that weakening of the sister chromatid cohesion around the chiasmata reduces the binding of the homologs in older oocytes, thus increasing their spatial separation.

### Lower quality oocytes are present in aged germline

In young adult worms, the diakinesis oocytes are stacked sequentially one after the other at the proximal end of the gonad. In aged worms, we found smaller oocytes which were aligned in multiple rows (Fig. 5A). Differential interference contrast (DIC) imaging confirmed that oocytes from aged worms were significantly smaller than those from young worms (Fig. 5B). Together with the presence of small oocytes, we also detected endomitotic nuclei starting at day 4. The number of endomitotic nuclei increased dramatically with age (30% at day 4, 87.5% at day 8, 96.3% at day 10; Fig. 1A). Endomitotic nuclei are oocytes that have bypassed the prophase I diakinesis arrest but that have failed to fully complete anaphase I and likely undergo endoreduplication instead of mitosis (McGee *et al.* 2012). Collectively, these results suggest that the arrest of oocytes in *C. elegans* can lead to various aberrations including lower quality oocytes, defective G2/M arrest, and reduced interhomolog cohesion.

**Figure 5:**
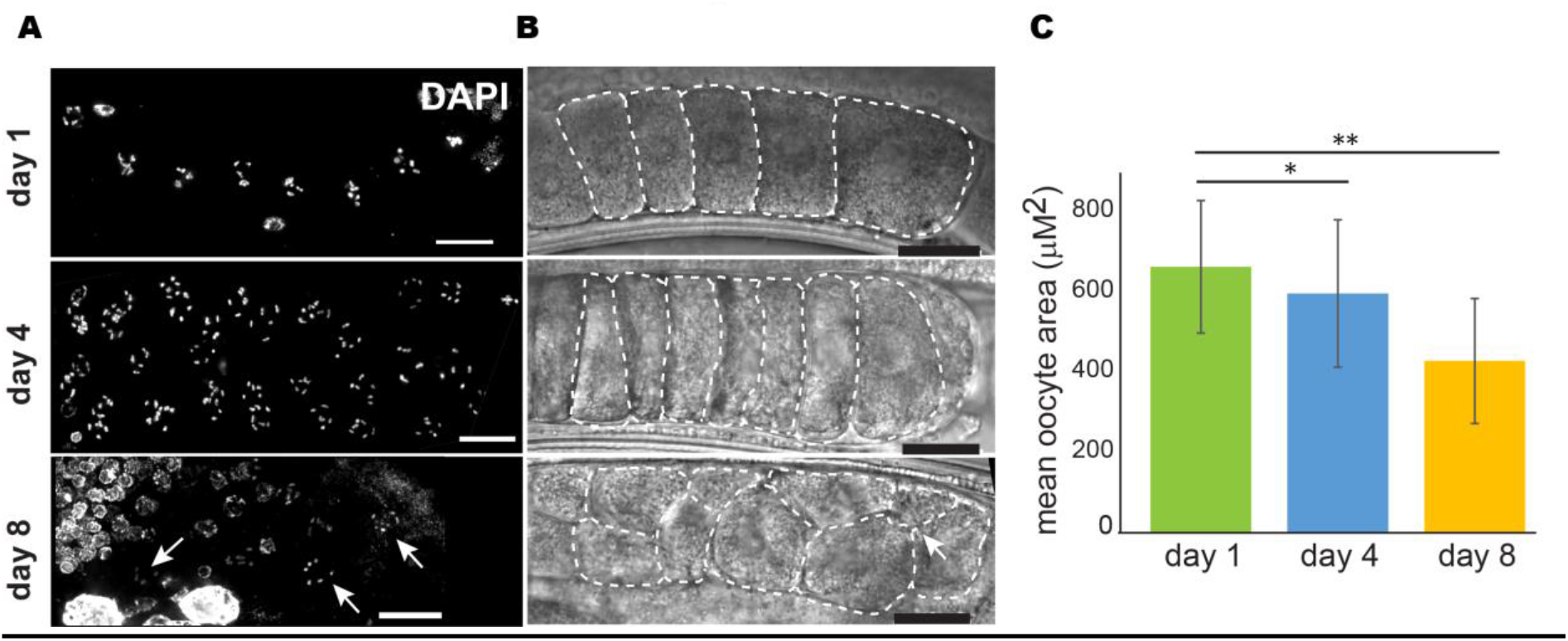
Oocyte get smaller with age. **A and B**. Proximal end of gonad arms at the indicated ages A) stained with DAPI and imaged with fluorescence microscopy and B) imaged with DIC. Arrows indicate pairs of diakinesis nuclei in the same row. White dashed lines indicate the circumstance of the mid plane of oocytes. Scale bar = 10 μM. **C**. Average area of oocytes at the different ages. Mann-Whitney *p* value: *<0.05, **<0.01. N = 6 gonads.

### MAPK signaling dynamics changes in aged gonads

We previously reported that a mutation in the *ogr-2* gene results in the development of small oocytes and endomitotic nuclei by day 3 post L4 (Achache *et al.* 2019). Similar phenotypes were observed in *lip-1* mutants (Hajnal & Berset 2002; Lee *et al.* 2006; Lin & Reinke 2008). The use of the *ogr-2* mutant strain is advantageous in germline studies, because unlike *lip-1,* its expression is limited to this tissue, and no somatic defects are observed in the deletion strain (Achache *et al.* 2019). Mutations in either of these genes lead to increased activation of MPK-1 in several regions of the gonads including mature oocytes at both day 1 and day 4 (Fig. S3) (Achache *et al.* 2019). The similarities between the phenotypes of old wild-type worms and day 3 mutant worms of these MPK-1 regulators, raise the hypothesis that MPK-1 activation changes with maternal age. Therefore, we stained the gonads of days 1, 4, 8, and 10 worms with an antibody directed against the phosphorylated (activated) form of MPK1 (dpMPK-1). As was previously demonstrated in young adult worms (Church *et al.* 1995; Lackner & Kim 1998; Kritikou *et al.* 2006; Lee *et al.* 2007; Arur *et al.* 2011; Yin *et al.* 2016; Narbonne *et al.* 2017), MPK-1 activation is restricted to two main regions of the gonad, the mid- to late-pachytene and the late diakinesis stages (Fig. 6). We observed that as the worm ages, MPK1 becomes ectopically activated in other regions of the gonad such as the proliferative zone, the leptotene/zygotene, and the diplotene as it is in young adult *lip-1* and *ogr-2* worms (Fig. 6) (Achache *et al.* 2019). The ectopic activation at diplotene was shown to lead to increased apoptosis in the germline of these strains (Rutkowski *et al.* 2011; Perrin *et al.* 2013). Importantly, unlike on day 1, at days 4, 8, and 10, the levels of dpMPK-1 staining at late diakinesis was similar to that at other stages (Fig. 6), probably due to depletion of sperm which activates MPK-1 through the major sperm protein MSP (Miller *et al.* 2001; Han *et al.* 2010). These results demonstrate that during germline aging, there is a change in dpMPK-1 staining dynamics along the gonad indicating that, like oogenesis progression, germline aging is linked to a change in MPK-1 activation.

**Figure 6:**
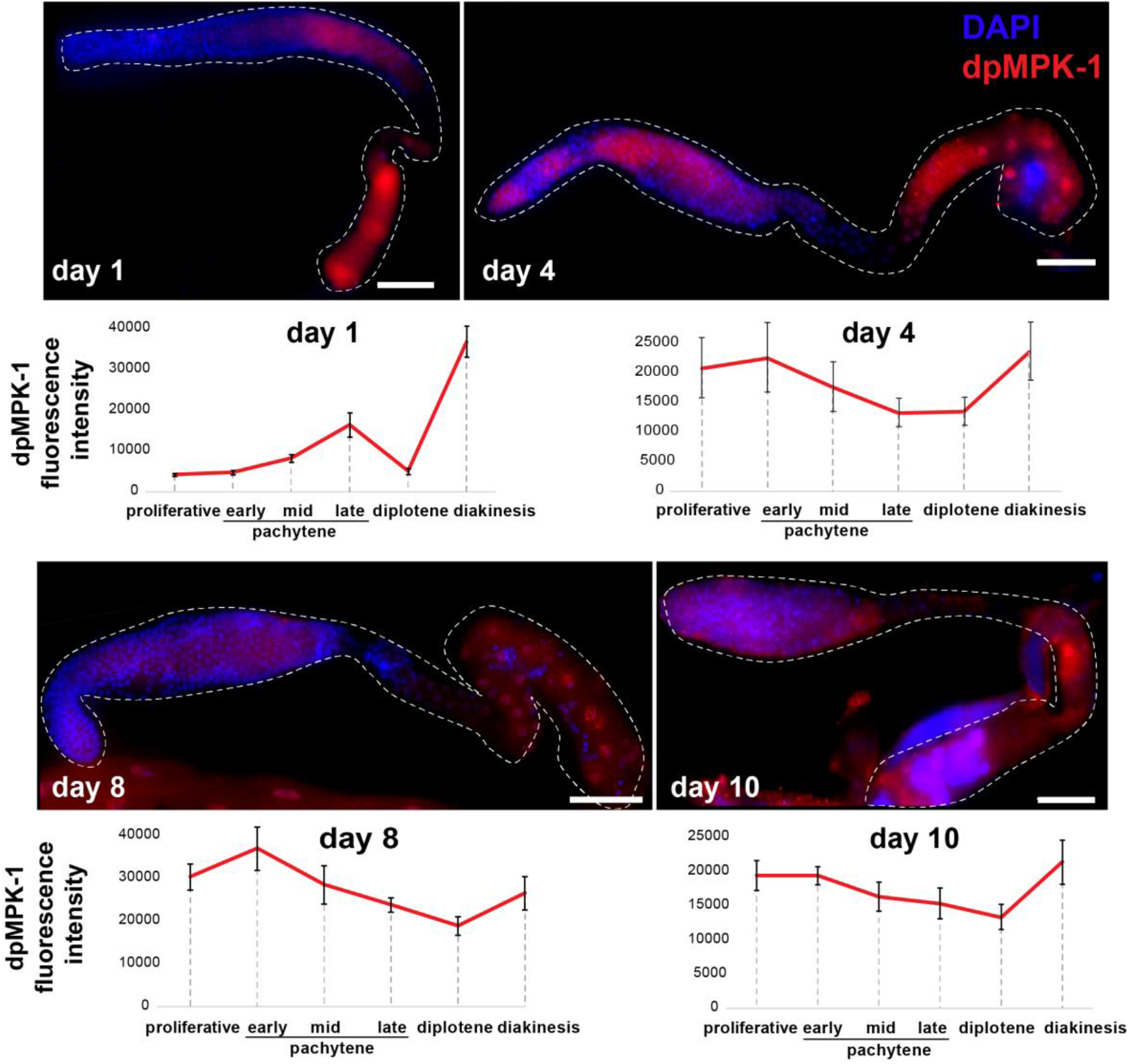
MPK-1 activation in the gonad changes during aging. Images of whole mount gonads from worms of the indicated ages stained for dpMPK-1 (red) and DAPI (blue). Average levels of dpMPK-1 signal are indicated below the images. N = 8 gonads for day 1, N = 8 for day 4, N = 5 for day 8, and N = 10 for day 10.

### Aged oocyte quality can be controlled by attenuation of MAPK signaling

Our findings that wild-type oocytes of aged worms have similar morphology to young oocytes with high levels of MPK-1 activation suggest that MAPK signaling influences oocyte quality throughout aging. To test this hypothesis, we compared the viability of aged fertilized oocytes with high and low levels of MAPK activation. This assay was established previously to evaluate oocyte quality (Andux & Ellis 2008). At 4 days post L4, wild-type, *lip-1,* and *ogr-2* worms were mated with young adult wild-type males. In these worms, the self-sperm was depleted approximately 24 hours before mating, so the oocytes were arrested for about one day. Between 8 and 12 hours after the introduction of males, adult worms (hermaphrodites and males) were removed, and the numbers of fertilized embryos were counted (Fig. 7A). This time window was chosen because we aimed to evaluate the quality of the embryos that originated from the stacked and aged oocytes only. After this window, the embryos laid could be meiocytes at the pachytene stage during the arrest. The hatched embryos were scored 24 and 48 hours after the removal of adult worms to assess embryonic viability. We found that the embryonic viability in mature wild-type mated worms was 67.1±18.5%. In contrast, the embryonic viability was significantly lower in mutants with higher MPK-1 activation: 44.7±28% for *ogr-2* and 22.6%±28% for *lip-1* (Fig. 7B). The difference between the two mutants could result from either different levels of MPK-1 activation (Fig. S3) or from somatic effects that exist in *lip-1* mutants but not in *ogr-2* mutants. We conclude that the quality of oocytes produced under conditions of MPK-1 overactivation is relative to the quality of oocytes produced from wild-type worms at the same aging step.

**Figure 7:**
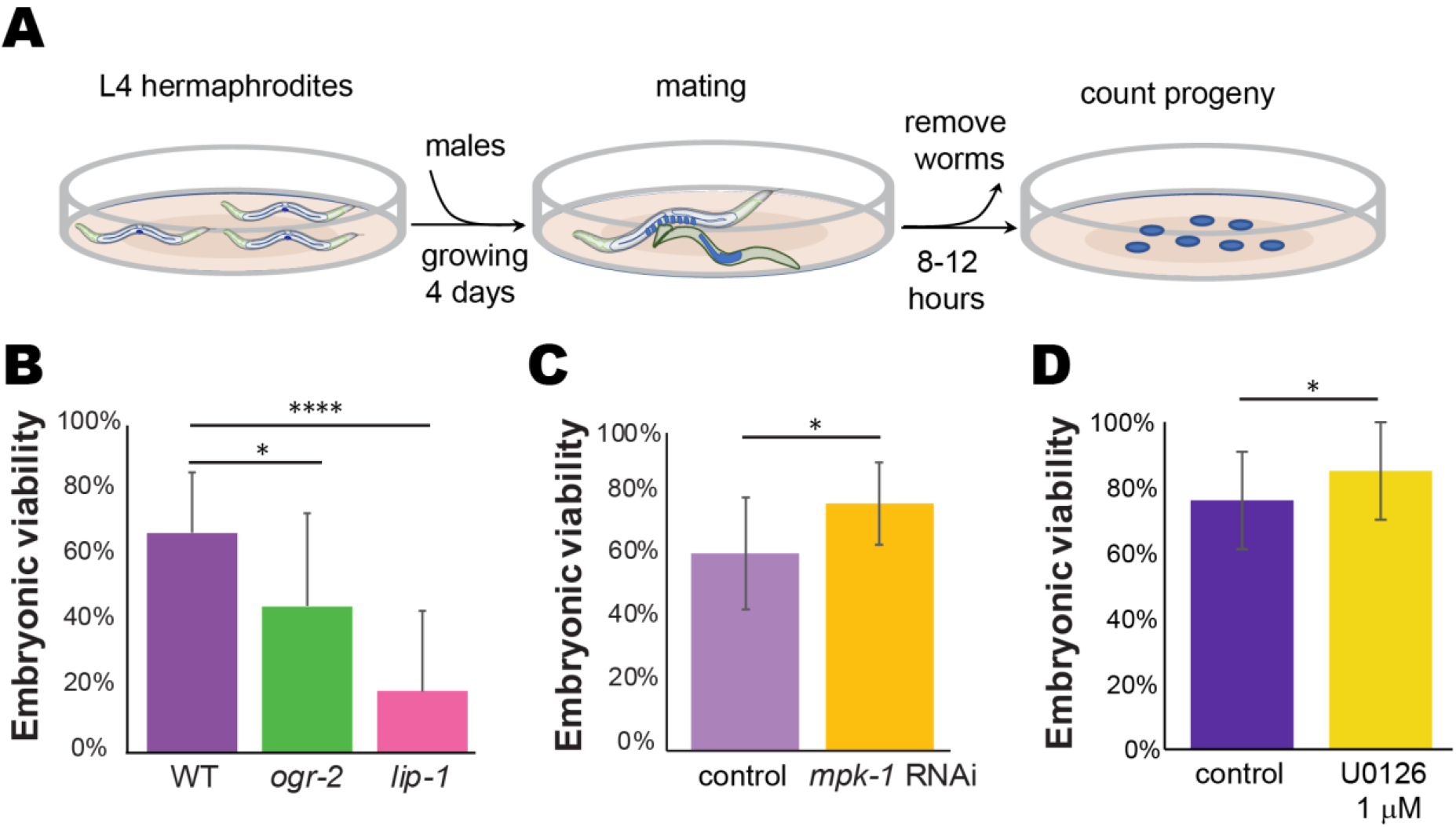
Viability of aged oocytes can be attenuated by MPK-1 activation. **A.** Illustration of the experimental design. **B-D.** Average embryonic viability of the first ~10 fertilized oocytes after the arrest in B) mutants that lead to over activation of MPK-1 in oocytes, C) in worms depleted of *mpk-1* using RNAi, and D) in worms that were kept in the presence an MPK-1 inhibitor during the arrest. Mann-Whitney *p* value: *<0.05, ***<0.001. Average of three different biological experiments. N = 20 worms.

Our hypothesis that the quality of arrested oocytes is determined by MPK-1 activation further predicts that reducing the levels of MPK-1 in the gonads will lead to an improvement in oocyte quality. MPK-1 mutants are either non-viable or lead to oogenesis arrest at late pachytene (Church *et al.* 1995), precluding mating experiments under similar conditions. We therefore used RNA silencing to partially reduce the levels of MPK-1 in the gonads. To specifically reduce the germline expression of MPK-1, we used the *rrf-1* strain in which somatic RNAi is reduced or abolished (Sijen *et al.* 2001). We found that the embryonic viability was significantly increased in oocytes with reduced *mpk-1* expression as compared to control RNAi (77.9±12.8% vs. 66.2±17.7%, respectively) (Fig. 7C). This result suggests that reducing MAPK signaling in arrested oocytes increases their quality and ability to successfully complete meiotic divisions or embryogenesis.

To test this, we used the specific ERK inhibitor U0126, which was shown to reduce MPK-1 activation in worm gonads (Morgan *et al.* 2010; Okuyama *et al.* 2010). Following self-sperm depletion, mature worms were moved to NGM plates containing the U0126 inhibitor. After 24 hours, the worms were returned to normal plates without the inhibitor, and the quality of the arrested oocytes was assessed by measuring the embryonic lethality of the first embryos that were laid after mating. We found that the quality of oocytes from worms exposed to the inhibitor was significantly increased compared to control worms (Fig. 7D). These results suggest that oocyte quality can be extended by reducing the MAPK signaling, even after oocytes are formed.

## Discussion

The classical model of aging defines it as the collective physiological processes that gradually decline, fail, and eventually lead to health deterioration and death over time (Lopez-Otin *et al.* 2013). The importance of focusing on oocyte aging, and the ways it can be delayed, is highlighted by the fact that oogenesis is one of the first processes to fail in humans and worms. Through careful analysis of oogonial processes along four critical points in the reproductive aging of *C. elegans,* we found an inherent signaling pathway that regulates oocyte aging. Our analyses indicate that several processes deteriorate with age. Indeed, we observed that the number of germ cells decreases, nuclei with a crescent shape morphology tend to disappear, the distance between the homologous chromosomes in mature oocytes increases, and there is a reduction in germ cell proliferation, crossover designation, and RAD-51 foci. In old worms the oocytes were smaller, and diakinetic nuclei were observed distally. In contrast, there were no differences in either synapsis or chiasmata formation between young and old worms.

When compared to previous work, one must keep in mind several differences. First, most previous analyses of oocytes at various aging stages (e.g., (Templeman & Murphy 2018)) used feminized mutants, whereas we strictly used the N2 wild-type strain. Thus, oocyte aging in our analyses started at the day 3 of adulthood, and not at day 1 as in the feminized strains. This difference may change the effects originating from somatic aging. Second, to reduce the number of days one has to move the worms to fresh plates due to the self-progeny, provide food, and avoid contamination, we kept the worms at 25 °C. Maintaining the worms at 25 °C allowed assessment of oocyte aging at day 4 when oocytes start to stack and arrest in contrast to day 5, at which this occurs in worms maintained at 20 °C. It is theoretically possible that differences in the aging dynamics exist between the two temperatures (Bilgir *et al.* 2013); however, our results are in agreement with previous publications (Hughes *et al.* 2007; Lim *et al.* 2008; Pickett *et al.* 2013; de la Guardia *et al.* 2016; Kocsisova *et al.* 2019) in terms of proliferation and oocyte morphology.

Some of our results can be explained by the halt in ovulation. This halt, which is mediated by the lack of sperm (McCarter *et al.* 1999), leads to oocyte stacking, and germ cell proliferation reduction, which together with ongoing apoptosis, leads to reduction in germ cell number. The disappearance of LZ nuclei can be the result of developmental progression of early meiocytes nuclei into pachytene, beyond the RAD-51 removal stage, without spatial movement, which explains the lower number of RAD-51 foci we found in aging gonads. The stacked oocytes gradually utilize their yolk and become smaller, whereas sister chromatid cohesion gradually weakens leading to homolog distancing and splayed COSA-1 signal.

Nevertheless, other results cannot be explained by this simplified ovulation model: the loss of oogenesis progression control, the increase in apoptosis in day 4 and appearance of endomitotic nuclei in aging oocytes. All these phenotypes have been connected in the past with aberrant MAPK signaling (Church *et al.* 1995; Lackner & Kim 1998; Hajnal & Berset 2002; Kritikou *et al.* 2006; Lee *et al.* 2007; Arur *et al.* 2011; Cha *et al.* 2012; Perrin *et al.* 2013; Yin *et al.* 2016; Narbonne *et al.* 2017; Achache *et al.* 2019), and indeed we found a dramatic change in the activation dynamics of MPK-1 in the gonads of aging worms. Together, this led us to suggest that MAPK signaling is a driver of the changes that occur in the aging germline. Several lines of evidence support this model. First, we found a dramatic alteration in the dynamics of MPK-1 activation during aging. Second, we found changes in proliferation, oogenesis staging, apoptosis, crossover designation, size of oocytes, and endomitosis, all previously shown to be controlled by MAPK (Lee *et al.* 2007). For example, we previously showed that ectopically high MPK-1 activation in the LZ region is associated with reduced LZ population (Achache *et al.* 2019), and Yin et al. showed that changes in local MPK-1 activation are associated with the appearance of diakinesis nuclei distally (Yin *et al.* 2016). Most importantly, here we showed that the period during which oocytes are of high quality can be extended or shortened by reducing or increasing, respectively, the level of MPK-1 activation using genetic and pharmacological tools. Taken together, we suggest that aging leads to a change in MAPK signaling in the gonad. Phosphorylation of downstream targets collectively lead to the different oogonial alternations. The change in MAPK signaling could be the result of germline intrinsic and/or extrinsic signals that come through the gonad sheath cells and/or as a result of lack of sperm (Miller *et al.* 2003; Govindan *et al.* 2006; Govindan *et al.* 2009; Li *et al.* 2012). A previous report showed that the IIS, known to control both longevity and germline aging, is inactivated in the absence of food leading to reduced MPK-1 activation and stalled oogenesis (Lopez *et al.* 2013). It is therefore possible that the IIS also works through MPK-1 to attenuate oogenesis progression and thus determine germline aging. In the future it will be interesting to test this hypothesis and to evaluate whether other signaling pathways influence germline aging (Qin & Hubbard 2015; Templeman & Murphy 2018) by attenuating MAPK signaling in the gonad. This will place MAPK as a link between the external signals and the internal processes of germline aging.

Our work, together with that of others (Hughes *et al.* 2007; Webster & Schuh 2017; Gruhn *et al.* 2019; Zielinska *et al.* 2019), highlight similarities in reproductive aging between worms and mammals. In both systems oocyte quality reduces with age, and the probability that oogenesis will be completed decreases with age. Changes in crossover were linked to age-related infertility in humans, and it also changes during worm reproductive aging ((Lim *et al.* 2008) and this work). Importantly, we found indications that aging leads to reduced sister chromatid cohesion in aged oocytes, as was also recently shown for mouse and human oocytes. If indeed the effects of MAPK on oocyte quality and maturation during aging are evolutionarily conserved, it will be critical to identify intrinsic downstream factors in the pathway that control oocyte aging in order to get deeper insights into human reproductive decline.

A landmark paper published by López-Otín et al. identified major features that constitute the hallmarks of aging (Lopez-Otin *et al.* 2013). Although this publication was focused on lifespan and healthspan, it defined the criteria for an aging hallmark: it appears during normal aging, its increase leads to accelerated aging and its reduction to slower aging. MAPK signaling can be regarded as a hallmark of oocyte aging by these criteria: First, MPK-1 activation levels in the oocytes change during normal aging. Second, when MPK-1 activation is increased as in *lip-1* and *ogr-2* oocytes, they age faster; mutant oocytes are smaller and more prone to pass the G2/M arrest, and the resulting embryo is less likely to complete embryogenesis. Most importantly, decreasing MPK-1 levels, by both RNAi and pharmacological inhibition, improved embryo viability. We conclude that our results indicate that MAPK is a major signaling pathway that attenuates oocyte aging.

## Methods and materials

### Strains and alleles

All strains were cultured under standard conditions at 25 °C except for mating experiments, *(lip-1, ogr-2,* RNAi and ERK inhibitor) which were conducted at 20° C (Brenner 1974). The N2 Bristol strain was used as the wild-type background. Worms were grown on NGM plates with *Escherichia coli* OP50 (Brenner 1974). The following mutations and chromosome rearrangements were used LGI: *rrf-1*(ok589), LGII: *ogr-2*(huj1), *meIs8* [pie-1p::GFP::cosa-1 + unc-119(+)], LGIV: *lip-1*(zh15), LGV: bcIs39 [Plim-7::ced-1::gfp+lin15(+)].

### Reproductive span analysis

To verify the effect of aging on the worm fertility, 400 L4 worms were placed on seeded NGM plates, transferred to new plates every 24 hours, and their embryos, non-fertilized oocytes, and hatched progeny were counted for 10 days at 25 °C.

### Gonad nuclei quantification

The numbers of nuclei at each meiotic stage, from the distal tip to the end of pachytene, were counted manually on DAPI-stained gonads as was previously described (Achache *et al.* 2019).

### Cytological analysis and immunostaining

Immunostaining of dissected gonads was carried out as described (Colaiacovo *et al.* 2003; Saito *et al.* 2009). Worms were permeabilized on Superfrost+ slides for 2 min with methanol at −20° and fixed for 30 min in 4% paraformaldehyde in PBS. After blocking with 1% BSA in PBS containing 0.1% Tween 20 (PBST) for 1 h at room temperature, slides were incubated with primary antibody for 1 h at room temperature. After incubation with fluorescent secondary antibody 1 h at room temperature, slides were DAPI stained for 10 min at 500 ng/ml, destained 1 h in PBST, and mounted with Vectashield (Vector Laboratories). The primary antibodies used were as follow: rabbit α-LAB-1 (1:200, (de Carvalho *et al.* 2008)), rabbit α-RAD-51 (1:10,000, SDIX), mouse α-MAPK-YT (1:500, M8159; Sigma), rabbit α-SYP-2 (1:200, a kind gift from S. Smolikove), rabbit α-pH3 (D5692, 1:1000; Sigma), and guinea pig α-HTP-3 (1:200, (Goodyer *et al.* 2008)). The secondary antibodies used were Cy2-donkey anti-rabbit, Cy3-donkey anti-guinea pig, Cy3-goat anti-rabbit, Cy3-goat anti-mouse (all used at 1:500 dilution; Jackson ImmunoResearch Laboratories).

### Imaging and microscopy

Images were acquired using the Olympus IX83 fluorescence microscope system (Olympus). Optical z-sections were collected at 0.30/0.60-μm increments with a Hamamatsu Orca Flash 4.0 v3 and CellSens Dimension imaging software (Olympus). Images were deconvolved using AutoQuant X3 (Media Cybernetics).

### Oocyte size measurement

Measurements were performed on whole worms mounted in M9 and visualized using DIC microscopy (Sulston & Horvitz 1977). Mid-oocytes plane areas were measured with ImageJ software.

### Quantitative analysis of germ cell apoptosis

Germ cell corpses were scored in adult hermaphrodites using CED-1::GFP as described (Zhou *et al.* 2001). Worms were transferred onto a drop of M9 on 1.5% agarose pads on slides and visualized. Statistical analyses were performed using the two-tailed Mann–Whitney *U*-test (95% C.I.).

### Quantification of immunofluorescence signals

Activated MPK-1 fluorescence intensity was quantified on raw images taken from whole-mounted gonads of wild-type worms at the different aging phases stained with an anti-dpMPK-1 antibody using the same experimental conditions and identical acquisition parameters. ImageJ software was used to measure the fluorescence intensity level throughout the entire length of the gonad.

### Time-course analysis for RAD-51 foci

The average RAD-51 foci number per nucleus was scored in each meiotic stage of the germline. Statistical comparisons between the different aging stages were performed using the two-tailed Mann–Whitney *U*-test (95% C.I.).

### Quantification of COSA-1 foci

For quantification of GFP::COSA-1 foci, nuclei that were in the last four-to-five rows of late pachytene and were completely contained within the image stack were analyzed. Foci were quantified manually from deconvolved 3D stacks.

### Quantification of the distance between the homologs

DAPI-stained images of the bivalents were recorded in z-stacks with vertical separation of Δz = *0.3μm* and horizontal pixel resolution of *Δx = Δy = 0.064μm.* To simplify the analysis, we selected bivalents in which the homolog interface (short arms) was parallel to the z axis (Fig. 4A). 3D fluorescence data were represented as a 3D matrix such that each voxel *(i,j,k)* has a measured fluorescence value *F(i,j,k).* First, we performed clustering-based segmentation in 3D to isolate the relevant bivalents. We then fitted the 3D fluorescence data to a Gaussian mixture model (GMM) with two Gaussians. Because GMM fitting operates on a point cloud rather than on a scalar intensity field, we represented the fluorescence matrix as a point cloud, in which the occurrence of each voxel’s coordinates is proportional to its fluorescence. Hence, the coordinates of each voxel *(i,j,k)* appeared in the point cloud round[0.1 · *F(i,j,k)]* times. The value of 0.1 was chosen to reduce the point-cloud’s size and speed up computation time; we verified that the results are unchanged when using a value of 1. GMM fitting was performed in Matlab^TM^ using the fitgmdist function with 20 replicates and 100 iterations per replicate. This procedure resulted in two Gaussians per bivalent, where each Gaussian in described by its mean (point in 3D) and 3 × 3 standard deviation matrix.

2D fluorescence density maps were obtained by summing all z-slices in a stack along the z direction. Such projections are shown in Fig. 4B, in which fluorescence is represented both by surface height and color code. The mean of each Gaussian was also projected onto the 2D map. The projection of each mean is very close to the two peaks in the 2D map due to the z-orientation of the bivalents in the raw data. The 2D distance *L* between the two bivalents was defined as the 2D distance between the 2D projected positions of the Gaussian means (Fig. 4B). The gap between the bivalent was further characterized by defining *H* as the fluorescence difference between the minimum value along *L* and the mean 2D fluorescence values at the two projected centers of the Gaussians (Fig. 4B). The overlap between the bivalents was quantified using the Jensen–Shannon Divergence (JSD), which measures the overlap between two probability distributions on a scale between 0 and 1 (Lin 1991; Endres & Schindelin 2003). A JSD value of 0 means the two distributions have no overlap, and a JSD value of 1 implies the two distributions are identical. To calculate JSD between the Gaussians fitted to the two bivalents, we first calculated the value of each Gaussian in each voxel and then normalized each Gaussian to 1, to make it a probability distribution. If *P_n_* and *Q_n_* are the values of the two Gaussians in the *n*’th voxel and 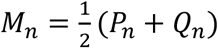, then the JSD is given by the sum over all voxels:

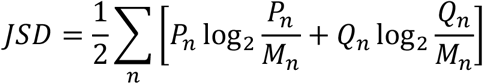

### Male generation

Young wild-type adult males were generated by crossing wild-type L4 hermaphrodites with wild-type males and growing for 3 days at 20 °C.

### Mating experiments

Twenty L4 worms were placed on seeded NGM plates and grown for 4 days at 20 °C. Hermaphrodites that have depleted their stock of sperm were mated with five males for 8-12 hours. This period corresponds to the time that takes the worms to lay between 10 to 15 eggs that originate from stacked oocytes. Hermaphrodites and males were then removed from the plate, and the embryos were counted immediately. Plates were scored for a second time after 24 hours. Embryos that had not hatched were marked as dead.

### Germline specific RNAi

Feeding RNAi experiments were performed at 20 °C as described (Govindan *et al.* 2006; Govindan *et al.* 2009). The control experiment was performed by feeding HT115 bacteria carrying the empty pL4440 vector. A feeding vector from the *C. elegans* RNAi collection (Source Biosciences) was used to deplete *mpk-1.* To study the germline-specific functions of MPK-1 in *C. elegans,* we used mutants of *rrf-1* strain, which encodes an RNA-directed RNA polymerase, to allow RNAi to be effective mostly in the germline (Sijen *et al.* 2001). Note that a previous report has shown that in some cases the RNAi in this strain also occurs in somatic tissues (Kumsta & Hansen 2012). Day 3 post L4, *rff-1* adult worms were placed on either *mpk-1* RNAi or control RNAi plates for 24 hours. They were then transferred to regular plates (to enable ovulation) and mated as described above.

### MPK-1 inhibitor assay

The MPK-1 inhibitor U0126 (1,4-diamino-2,3-dicyano-1,4-bis[2-aminophenylthio] butadiene monoethanolate) was purchased from Sigma. To test whether U0126 influenced oocyte quality, we treated day 4 adults for 24 h with either DMSO or 1 μM U0126. Adults were transferred to plates with 1 μl of 10 mM U0126 or DMSO. After 24 h, worms were transferred to regular NGM plates and mated with males. We then scored embryo production after 8-12 hours of mating as readout of oocyte quality.

## Acknowledgments

We thank the Caenorhabditis Genetics Center for kindly providing strains. We thank Sarit Smolikove for the SYP-2 antibody. This work was supported by Israel Science Foundation (grant numbers 1283/15, 2090/15) to Y.B.T. and the Faculty Fellowship of the Azrieli Fellows Program to T.B.

## Availability of data and materials

Strains and plasmids are available upon request.

## Authors’ contributions

RF and HA performed experiments and analyzed data. TB and NL devised and performed computational and biophysical analysis. HA and YBT designed experiments and wrote the manuscript.

## Supplementary Figures

**Figure S1:**
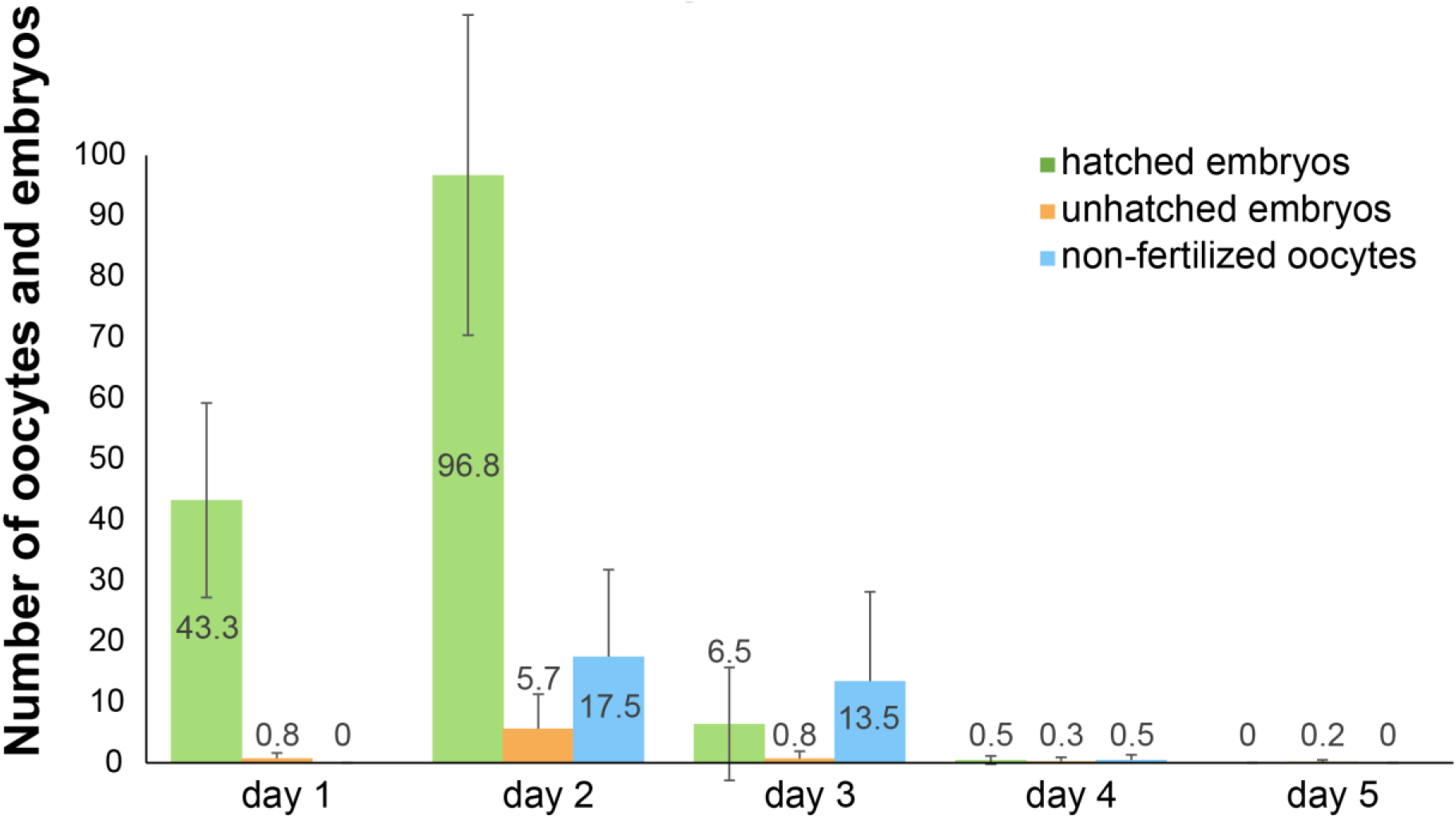
Ovulation ceases at day four of adulthood. Average number of oocytes, hatched and unhatched embryos detected at different days of adulthood. N = 20 worms.

**Figure S2:**
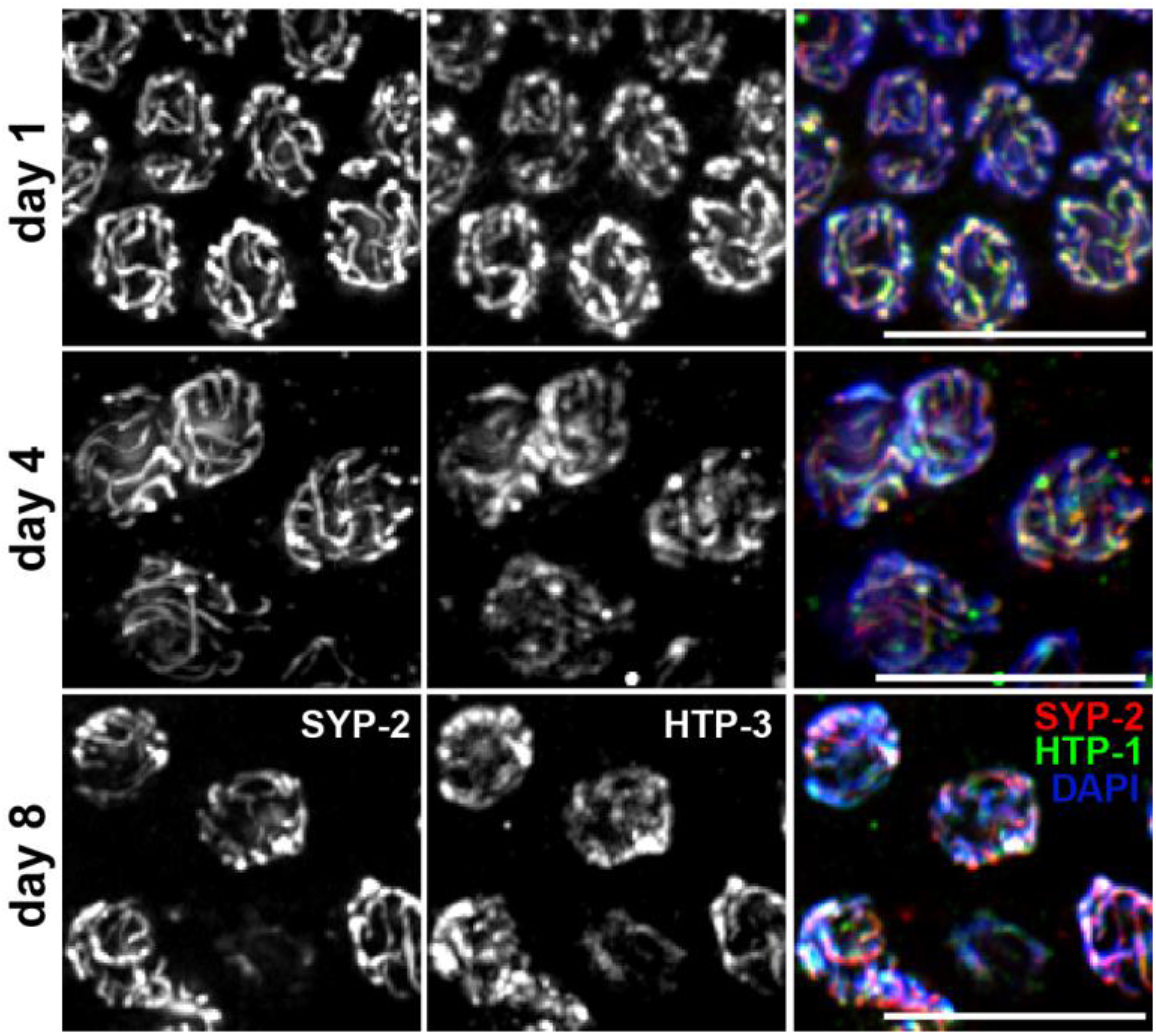
The synaptonemal complex is properly formed in aged germlines. SYP-2 (red), HTP-3 (green), and DAPI (blue) stained images of mid-pachytene nuclei at the indicated ages. Scale bar = 10 μM.

**Figure S3:**
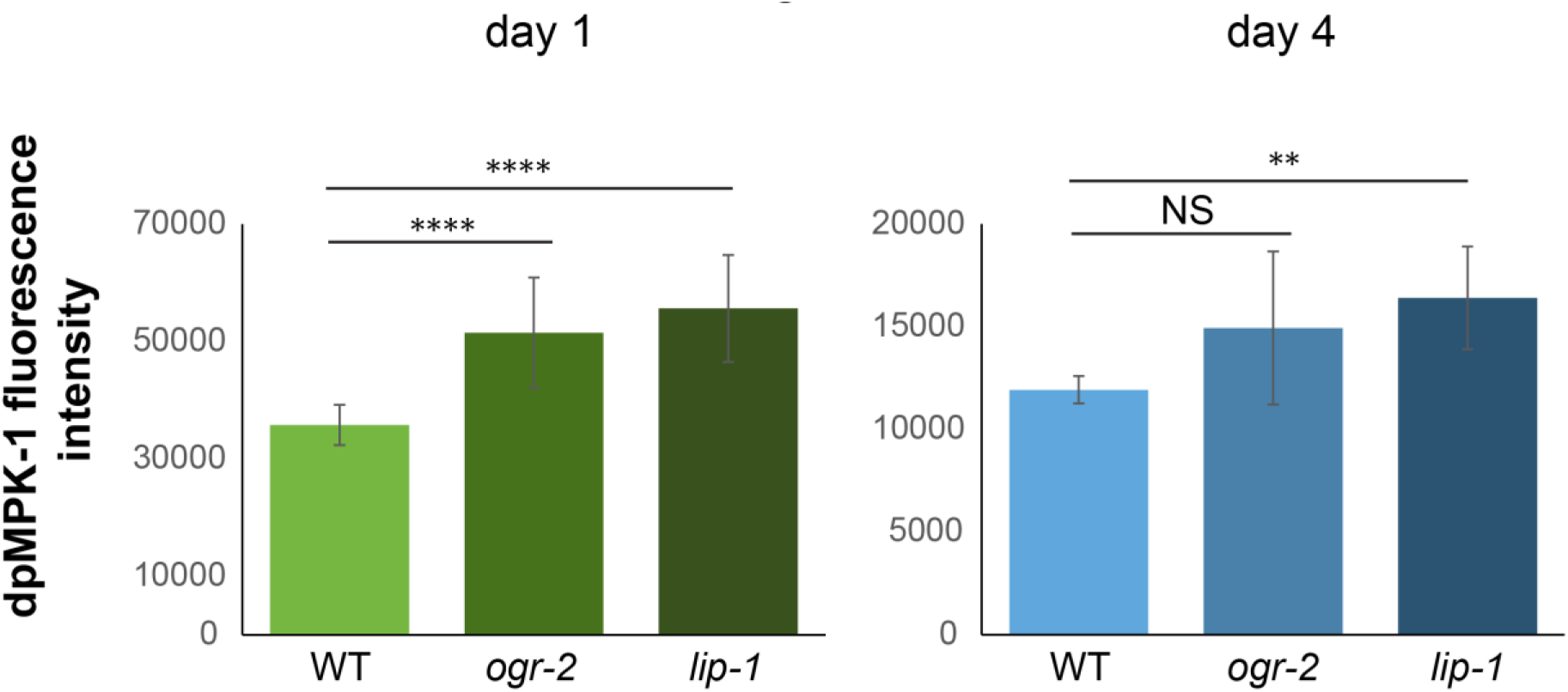
MPK-1 activation is higher in oocytes with *ogr-2* and *lip-1* mutations. Average fluorescence intensity of staining for dpMPK-1 in mature oocytes of the indicted genotypes at day 1 and day 4 of adulthood. Mann-Whitney *p* value: NS – not significant, **<0.01, ****<0.0001. N value =12 gonads.

